# Automated quantification of vomeronasal glomeruli number, size, and color composition after immunofluorescent staining

**DOI:** 10.1101/2021.06.24.449743

**Authors:** Shahab Bahreini Jangjoo, Jennifer M Lin, Farhood Etaati, Sydney Fearnley, Jean-François Cloutier, Alexander Khmaladze, Paolo E. Forni

## Abstract

Glomeruli are neuropil rich regions of the main or accessory olfactory bulbs where the axons of olfactory or vomeronasal neurons and dendrites of mitral/tufted cells form synaptic connections. In the main olfactory system olfactory sensory neurons (OSNs) expressing the same receptor innervate one or two glomeruli. However, in the accessory olfactory system, vomeronasal sensory neurons (VSNs) expressing the same receptor can innervate up to 30 different glomeruli in the accessory olfactory bulb (AOB).

Genetic mutation disrupting genes with a role in defining the identity/diversity of olfactory and vomeronasal neurons can alter number and size of glomeruli. Interestingly, two cell surface molecules, Kirrel2 and Kirrel3, have been indicated to play a critical role in the organization of axons into glomeruli in the AOB.

Being able to quantify differences in glomeruli features such as number, size or immunoreactivity for specific markers is an important experimental approach to validate the role of specific genes in controlling neuronal connectivity and circuit formation in control or mutant animals. Since the manual recognition and quantification of glomeruli on digital images is a challenging and time-consuming task, we generated a program in Python able to identify glomeruli in digital images and quantify their properties, such as size, number, and pixel intensity. Validation of our program indicates that our script is a fast and suitable tool for high throughput quantification of glomerular features of mouse lines with different genetic makeup.

## INTRODUCTION

An important aspect in neuroscience is to identify genes responsible for synaptic target specificity. Guidance cues, surface molecules, and cell adhesion molecules are key players in axon guidance and synapse formation across different types of neurons (Cloutier *et al.* 2002; Dean *et al.* 2003; Martin *et al.* 2015). Vomeronasal sensory neurons (VSNs) located in the vomeronasal organ (VNO) express vomeronasal receptors (VRs) which detect certain chemosignals and transduce that signal to the accessory olfactory bulb (AOB) in the brain.

Most of the neurons of the vomeronasal epithelium (VNE) belong to two main populations which detect different types of pheromones and play different roles in elicited social responses. VSNs located in the apical region of the vomeronasal epithelium express members of the vomeronasal receptor type 1 (V1rs) family which transduce their signal through the Gαi2 G-protein subunit, express the transcription factor (TF) Meis2 and the guidance cue receptor Nrp2 (Dulac and Axel 1995; Enomoto *et al.* 2011). These neurons project their axons to the anterior portion of the AOB. VSNs located in the basal region of the VNE express vomeronasal type 2 receptors (V2rs) (Silvotti *et al.* 2011), transduce their signal through the Gαo G-protein subunit, express the TF AP-2ɛ(Enomoto *et al.* 2011; Lin *et al.* 2018), and the guidance cue receptor Robo2. These neurons project their axons to the posterior AOB (Figure 1). A third subtype of VSNs express Formyl peptide receptors (Fprs)(Bufe *et al.* 2012). Neurons expressing mFpr-rs1 also express Gαo whereas Gαi2 expressing neurons can express mFpr-rs3, mFpr-rs4, mFpr-rs6, and mFpr-rs7 (Bufe *et al.* 2012; Riviere *et al.* 2009). Proper differentiation, organization, and signaling in VSNs are necessary for their individual roles in integrating external cues and translating them into behaviors (Brignall and Cloutier 2015; Chamero *et al.* 2012).

**Figure 1.**
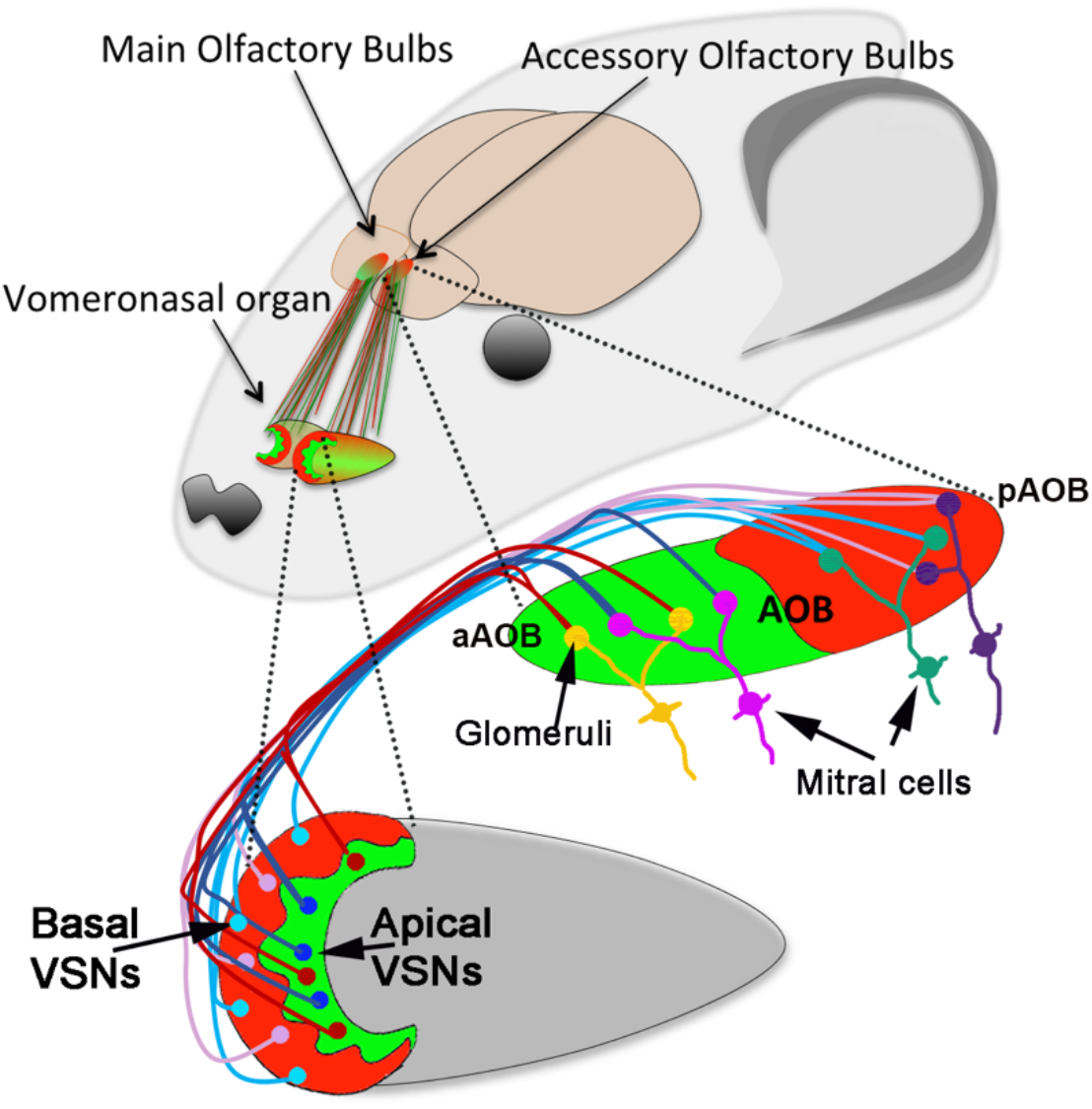
Cartoon of mouse head. The VNO projects to the accessory olfactory bulb (AOB) (red, green). Schematic of the vomeronasal epithelium: VSNs in the apical region (green) project to the anterior AOB (a-AOB, green) while VSNs in the basal areas (red) project to the posterior AOB (p-AOB, red). VSNs, based on the receptor that they express (indicated by different colors: cyan, pink, blue and red) connect with specific mitral cells in the AOB (yellow, magenta, green, and purple) forming distinct glomeruli in the AOB.

Glomeruli are neuropil rich regions of the olfactory bulbs where the axons of olfactory or vomeronasal neurons and dendrites of mitral/tufted cells form synaptic connections. The organization of axons and the glomeruli they innervate largely relies on the type of olfactory/vomeronasal receptor gene expressed by individual neurons. The main olfactory sensory neurons (OSNs) expressing the same receptor innervate one or two glomeruli. However, in the accessory olfactory system, vomeronasal sensory neurons (VSNs) expressing the same receptor can innervate up to 30 different glomeruli in the accessory olfactory bulb (AOB)(Chamero *et al.* 2012).

The fasciculation of the vomeronasal axons and formation of synaptic connections with their targets in the brain are mediated by the guidance molecules, the vomeronasal receptor(s) expressed, and by complex expression of cell surface molecules (Brignall and Cloutier 2015). The Kirrel family of cell surface molecules have been previously shown to play a relevant role in controlling axon coalescence in the olfactory and vomeronasal systems (Prince *et al.* 2013; Serizawa *et al.* 2006; Vaddadi *et al.* 2019). Combinatorial expression of Kirrel2 and Kirrel3 in VSNs assist in the formation and organization of glomeruli in the AOB. Notably, neurons projecting to the anterior portion of the AOB mostly express Kirrel2 while the basal neurons, projecting to the posterior AOB are mostly positive for Kirrel3 (Figure 3B)(Brignall and Cloutier 2015; Prince *et al.* 2013).

When experimentally analyzed, glomeruli are usually hand traced on digital images to generate regions of interest (ROIs) which can then be analyzed for number, size, and color composition or intensity using image analysis software, such as ImageJ (Naik *et al.* 2020; Prince *et al.* 2013). On histological sections glomeruli are commonly identified as OMP-positive and/or VGluT2 positive structures surrounded by a non-innervated region (Prince *et al.* 2013).

Notably, human made identification of glomeruli can be inconsistent, as it is dependent on the investigators’ ability to recognize complex morphological structures. These factors can compromise consistency in measurement when images are analyzed by different researchers. Additionally, this kind of analysis is usually tedious and time consuming. In order to facilitate this process and increase consistency, we generated an image processing program in Python that can be used to quantify glomeruli size, number, and immunostaining intensity.

The program employs OpenCV, which is an open source image processing library (Bradski 2000) that has been previously used in biological applications to identify patterns in an image on digital images (Cosentino *et al.* 2015; Dominguez *et al.* 2017; Lutnick *et al.* 2019; Obando *et al.* 2018; Uchida 2013). Our software separates multichannel images into single color images, uses adaptive thresholding to identify patterns, and quantifies features such as area, number of objects, and intensity for the selected channels.

We validated our bioinformatic tool and tested various parameters on samples of previously published wild-type controls and mutant animals carrying genetic mutations affecting glomeruli protein expression, size, and number (Prince *et al.* 2013). By comparing human and automated identification of glomeruli we noticed obvious differences in what would be considered one or multiple glomeruli. However, when normalized, the discrepancies between manual and automated quantifications did not compromise the detection of significant differences between genotypes. Our data suggest that our script can be used as a suitable tool to perform high-throughput analysis of glomeruli features and identify differences between mice with different genetic makeup.

## MATERIALS AND METHODS

### Manual Quantification of Glomeruli Structure and Formation

Selection of glomeruli were performed on 20μm parasagittal cryosections of adult mice immunostained with markers expressed by the glomeruli in the AOB and imaged through confocal microscopy. Glomeruli in the anterior and posterior regions of the AOB were defined as discrete regions of OMP+ or VGluT2+ axonal endings with similar composition of Kirrel2 and Kirrel3 expression that differ from neighboring regions. Anterior and posterior regions of the AOB were identified based on differential staining against Nrp2, Robo2, and BS-Lectin (Antibodies and concentrations used can be found in Key Resources Table). Manual quantifications for glomerular size, number per section, and immunoreactivity were performed on confocal images using FIJI 2.1.0.

### Image Preparation for Automated Analysis

Confocal images were imported into FIJI 2.1.0, where the relevant Z-stacks were combined using Maximum Intensity Z-projection. The individual ROIs were generated by identifying the anterior and posterior glomerular regions of the AOB through morphological and histological means. The area outside of the desired region to quantify was cropped out to limit quantification of irrelevant regions. All images were saved as TIFF files to preserve the individual channels.

### Development of Image Analysis Program

This program is written in Python 3 generation programing language. The images are subjected to thresholding, which is done by ADAPTIVE_THRESH_GAUSSIAN_C method in OpenCV, to convert images into binary. Also, the constant subtraction value for chosen Gaussian thresholding set to 4 for all images. This specific method requires a set block size, which determines the number of pixels surrounding a given pixel that should be considered to set the threshold limit. The threshold for each pixel is then set by comparing the intensity the pixel intensity to the average intensity of all the pixels in the block. The object identification was performed using three different squared block sizes to check the program output sensitivity for this parameter. Smaller block sizes consider smaller area in a pixel proximity and results in smaller identified objects on average.

For glomeruli identification, the program uses OpenCV library and its “contours” tool. RETR_EXTERNAL flag is implemented to pick the most outer region of identified objects and ignore all inside complexity such as holes. It is also known as “parent contours” in OpenCV. Another restriction for detected patterns is size limit. This limit set a minimum threshold to ignore all miniscule contours. All processes were done for six different minimum size limits to check how this parameter affects the ability to detect phenotypic differences between control and wildtype animals.

In this paper we only used Block size and minimum size limit to identify glomeruli, however, other functions that may assist in glomeruli detection and/or limit the identification of objects that are not glomeruli (e.g., nerve fibers) such as morphology filter tools (i.e., circular filter, ellipse filter, ratio filter), histograms, and selection filters are still in development.

Instructions for installation, program requirements, and access to the code are accessible as supplementary data and at https://github.com/ForniLab/Glomeruli-Analyzer.

### Experimental Design and Statistical Analysis

Data from the automated analysis were generated for area (# of pixels) and average intensity level per pixel. Human data were generated for area as <u>μm^2^ and intensity as corrected total fluorescence level (CTF) per μm^2^</u>. Data from manual and automated analysis were exported to Microsoft Excel. For each experiment we analyzed at least 3 samples per genotype/condition. The values (size, number of glomeruli, intensity) for each sample were averaged as one datapoint. Averaged data were imported and analyzed using GraphPad Prism9. **Comparison of manual and automated values were performed on normalized values to the average of each dataset. All data were collected from mice kept under similar housing conditions, in transparent cages on a normal 12 hr. light/dark cycle. Tissue collected from males and females in the same genotype/treatment group were combined; ages analyzed are indicated in text and figures. Sample sizes and p-values are indicated as single points in each graph and/or in figure legends. The data are presented as mean ± SEM. Two-tailed, unpaired t-test were used for all statistical analyses, and calculated p-values <0.05 were considered statistically significant.

### Automated Analysis of Images Using Gaussian Adaptive Thresholding

The human identification of glomeruli is based on prior knowledge or an expectation of a probable shape. Therefore, the experience and personal preferences can affect manual selection of glomeruli (Beck and Kastner 2009).

Our goal when developing this program was to identify “objects”/regions of interest (ROIs) based on continuous regions of color composition and intensity.

As a first step, the software separates multichannel (Figure 2A) images into individual channels (Figure 2 B). The target colors and staining are supposed to represent glomeruli areas as illuminated regions (Figure 2C). To identify distinct patterns, based on relative intensity, we used adaptive Gaussian threshold (Otsu 1979) (Figure 2D). This thresholding technique is based on the relative intensity of neighboring regions. The threshold for each pixel is set by comparing the intensity of each pixel to the average intensity of the pixels in the surrounding area. The surrounding area is a square region is referred as “block” and the number of pixels included in this area is referred to in this paper as “block size.” If the average intensity of all the pixels in the block is greater than the intensity of the pixel, the pixel is discarded.

**Figure 2.**
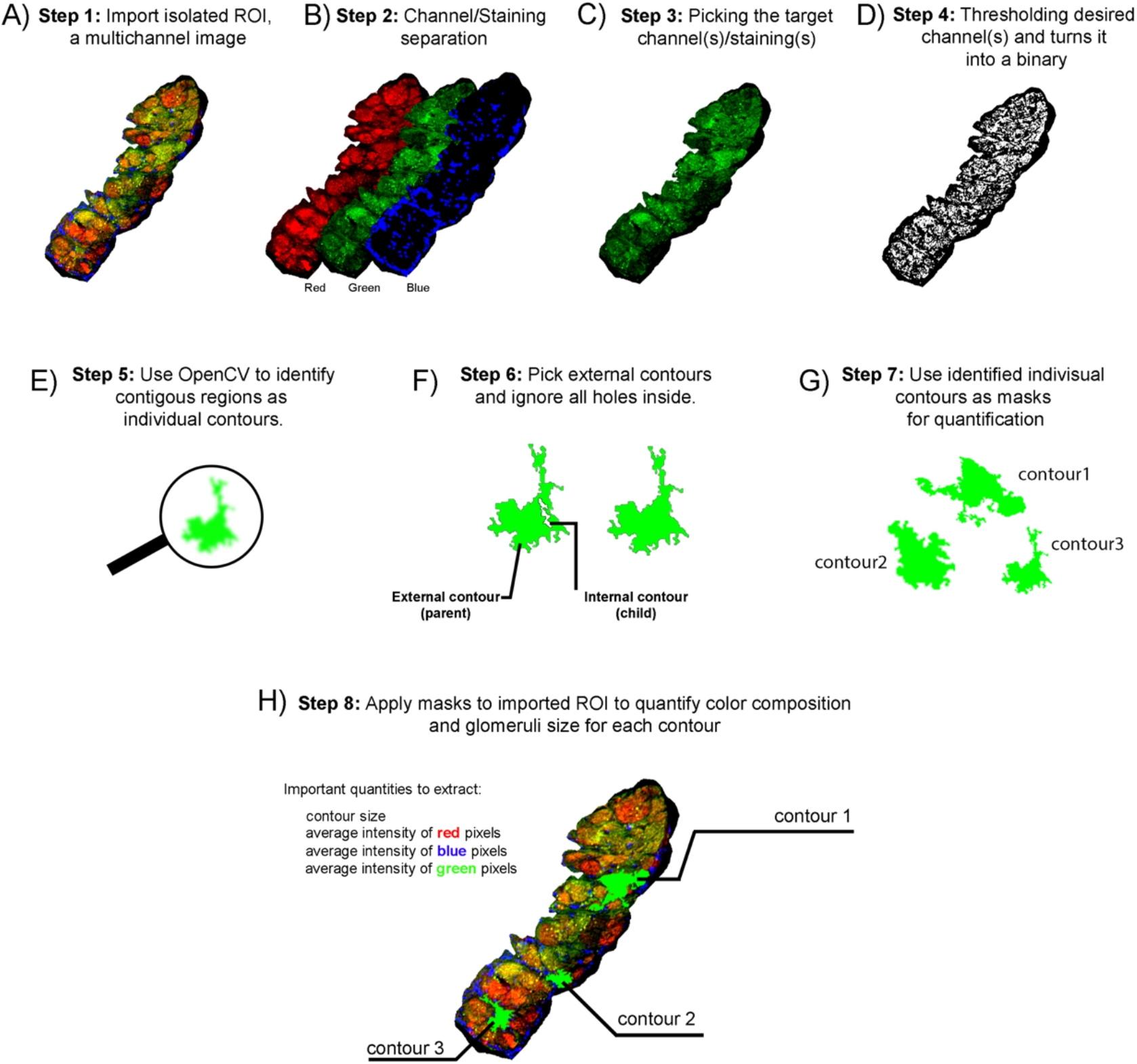
Diagram showing how the program identifies putative glomeruli in an ROI. A) The multichannel image of the anterior or posterior AOB (a multicolor image) is imported. In this RGB image the channels are stainings against BS-Lectin (red), Olfactory marker protein (OMP, green), and Hoescht (blue) counterstain, respectively. B) Split channels to isolate the separate staining. C) Choose the channel with the specific marker expressed by the glomeruli. D) Gaussian adaptive threshold is applied to the selected channel. E) The binary image is fed into the contour OpenCV algorithm to find all available objects and record all the identified objects and their locations. F) Use pattern recognition to select only the outermost/external contours (parent contours) and ignore all holes inside the identified objects/contours (child contours). G) Use the previously identified patterns/contours to create masks and crop its location from originally imported image and analyze the region. H) Using these masks, the program finds average intensity for each channel (e.g., red, green, and blue) for each of the previously identified contours from the original image. The program exports data for the number of identified contours in the entire image, average color intensities per contour, and average contour size (# of pixels) for the imported image.

To identify glomeruli, the binary channels are processed by OpenCV library contour finding function to seek all possible external contours (Figure 2E). The contour size is defined as all pixels that are enclosed by the contour border. Contours can be as small as 1 pixel in size. To discriminate between objects of interest and noise we set minimum size limits for the identified contours (Figure 2F). Throughout this paper, we set different block sizes and minimum size limits to test the influence of these parameters in identifying distinct glomeruli (see Figure 4). This program extracts average pixel intensity and the size per contour and records their numbers in aAOB and pAOB by creating masks out of identified contours (Figure 2G).

## RESULTS AND DISCUSSION

### Detection and quantifications of glomeruli size and immunoreactivity in the anterior and posterior AOB of WT animals

We tested our script on histological sections of the AOB of adult wild type mice, where glomeruli of the anterior and posterior AOB were previously manually identified and traced based on immunoreactivity against Robo2, Nrp2 (Figure 3A), and anti-VGluT2 (Figure 3A’), or Kirrel2, Kirrel3 (Figure 3B), and anti-VGluT2 (Figure 3B’). The manually made ROIs for the glomeruli were used to quantify the average glomerular size (μm^2^), and average number of glomeruli in aAOB and pAOB per section. The same ROIs were used to quantify and the average immunoreactivity for Kirrel2 and Kirrel3 for the glomeruli in each region.

**Figure 3.**
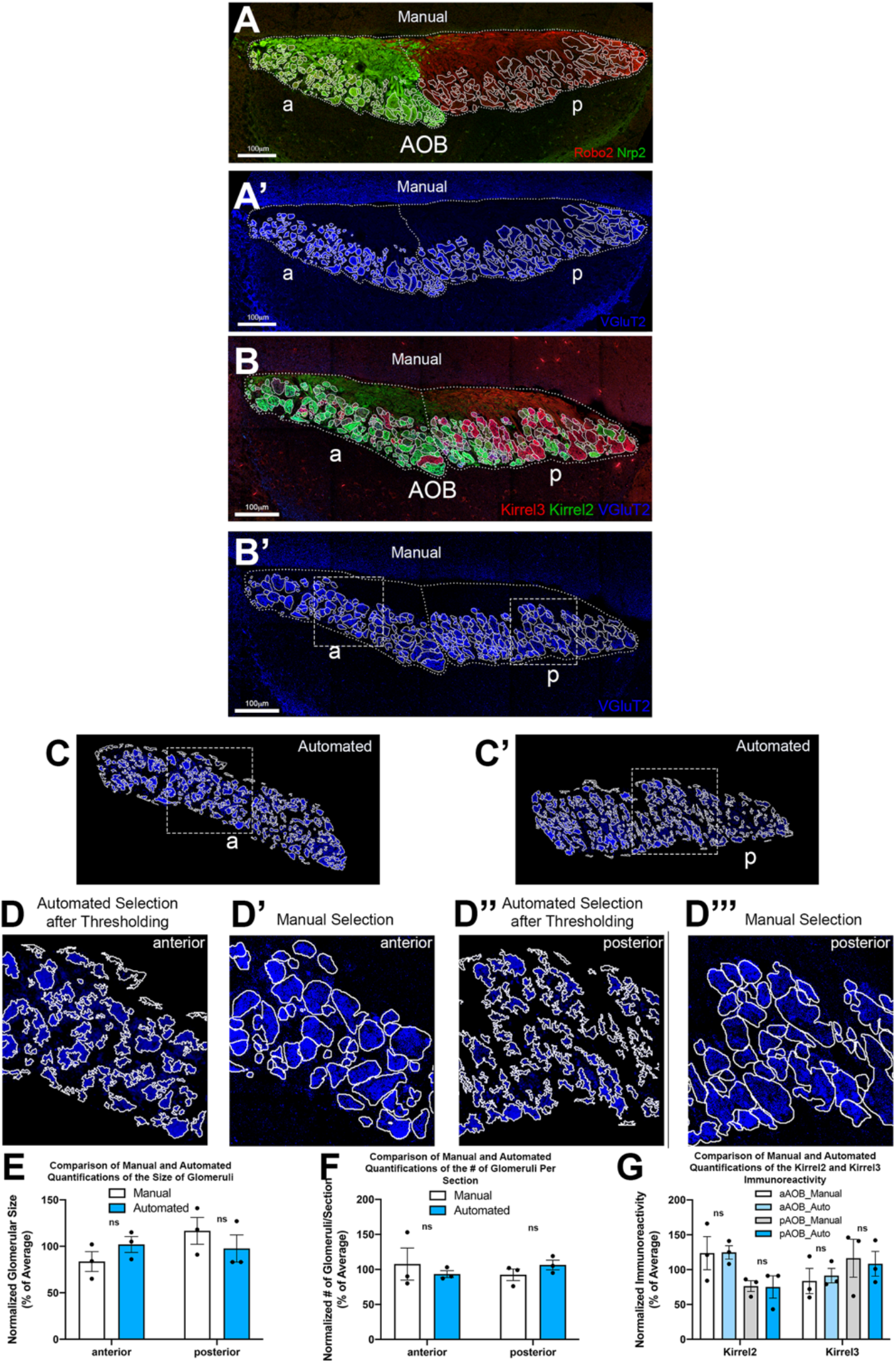
Comparison of manual and automated selections of glomeruli in the anterior and posterior regions of the accessory olfactory bulb (AOB) in an adult wildtype mouse. The VGluT2 (blue) positive glomeruli (Block size 101 and limit size of 40 has been set for this analysis.) (A’, B’) in the AOB were manually identified and traced throughout the anterior (a) and posterior (p) regions. Identification of anterior and posterior AOB regions can be done based on morphology and immunohistochemical markers. A-A’) Immunostainings against Nrp2 (green) and Robo2 (red) highlight the aAOB and pAOB, respectively. B-B’) Glomeruli throughout the AOB have varied composition of the cell surface molecules Kirrel2 (green) and Kirrel3 (red). C) Automated identification of glomeruli in anterior (C) and posterior (C’) regions based on VGluT2 (blue) immunostaining. D-D’’’) High magnification comparisons of automated selections after thresholding and manual selections of the same regions in anterior (D-D’) and posterior (D’’-D’’’) Dashed rectangle regions (B’, C, C’) indicate regions used for magnified comparisons for automated and manually selected glomeruli/ E-G) Comparison of the normalized values from manual and automated glomeruli selections showed no statistically significant differences for the E) average size of glomeruli, F) # of glomeruli/section, and G) immunoreactivity of Kirrel2 and Kirrel3 in anterior and posterior regions of an adult wild-type AOB.

For automated quantification, our script was able to identify glomeruli in the aAOB (Figure 3C) and pAOB (Figure 3C’). Dashed rectangles show same regions in the anterior AOB (Figure 3B’, C) and posterior AOB (Figure 3B’, C’). The magnified selections of the anterior are shown in Figure 3D and D’ and the magnified selection for the posterior are shown in Figure 3D’’ and D’’’. Figure 3D,D’’ are resulting glomeruli identified by the program and Figure 3D’, D’’’ are the manually identified glomeruli. Here, the blue channel (VGluT2) was used, and adaptive threshold block size was set to 101 and all contours below 40 pixels in size were discarded. The presumptive glomeruli areas identified by the program after thresholding appeared obviously smaller and surrounded by sharper edges compared to what can be identified by humans on non-naive images (Figure 3D-D’”). The automated selections yielded glomerular area (# of pixels), number of glomeruli per section, and immunofluorescence. To compare automated and manual methods, the values for glomerular area (Figure 3E), number per section (Figure 3F), and differential immunoreactivity for Kirrel2 and Kirrel3 (Figure 3G) in anterior and posterior AOBs were normalized to the average of each dataset. The automated quantifications did not result to be significantly different from the human made ones, suggesting that our script could be further tested to analyze different parameters of genetically modified mouse models.

### A comparison of human and automated quantification of Kirrel3 mutants vs controls

To understand if this program was capable of detecting the differences in glomerular organization between mice carrying different genetic mutations, we analyzed the same image sets used to generate the previously published data for WT controls (Figure 4A) and Kirrel3 KO (Figure 4B) mutant mice where the glomeruli in the pAOB were estimated to be approximately twice as large and about half as many glomeruli in the pAOB, highlighted by BS-Lectin, when compared to WT controls (Prince *et al.* 2013). The confocal images were divided into separate sets of images containing either anterior or posterior AOBs for each animal and condition before being put through the automated analysis (as illustrated in Figure 3C, C’).

**Figure 4.**
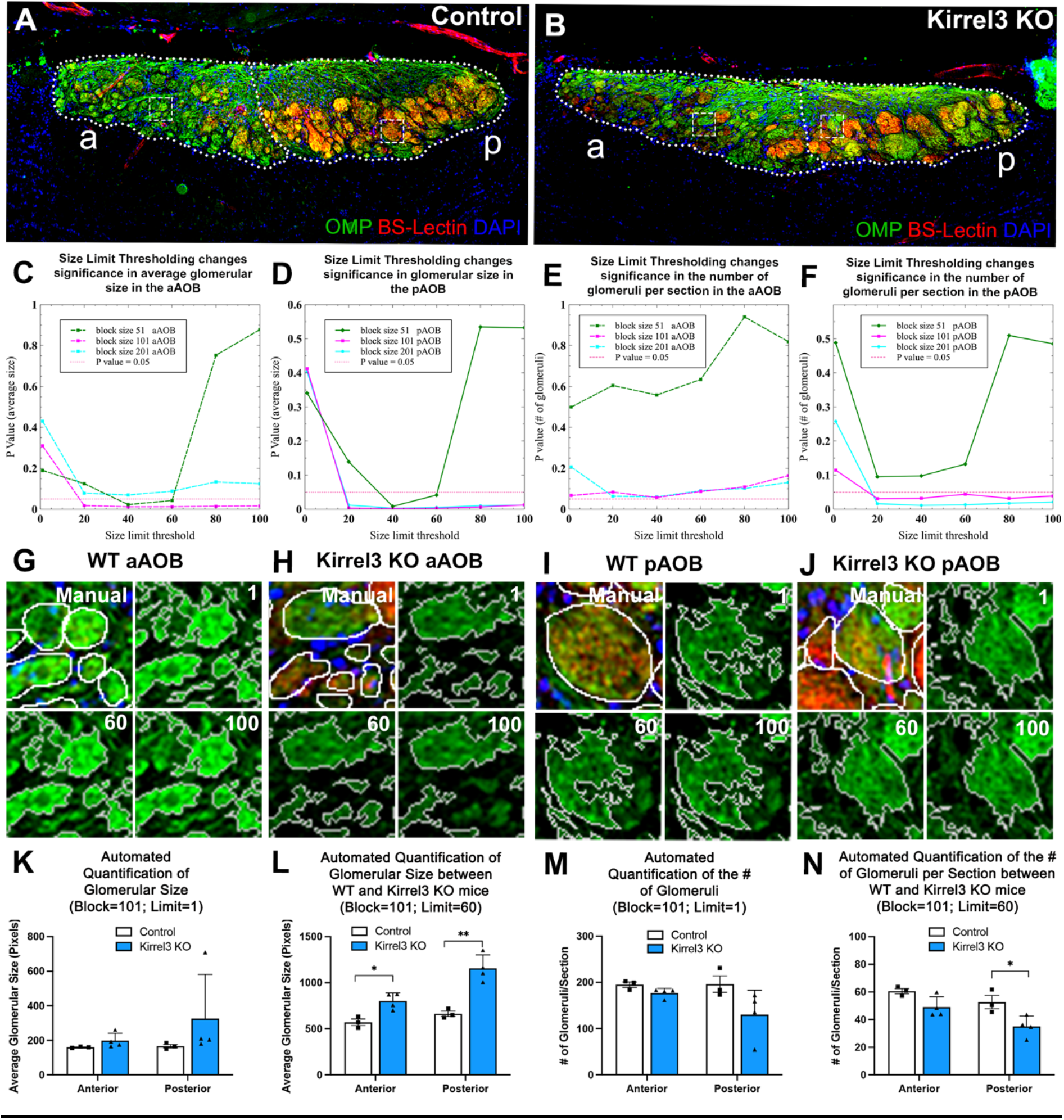
The automated selection and analyses of glomeruli in Kirrel3 mutants and WT controls. A-B) The olfactory marker protein (OMP, green) immunostaining and the BS-Lectin (red) staining with a Hoescht (blue) counterstain on the AOBs of adult WT control (A) and Kirrel3 KO mutant (B). OMP is present throughout the fibers of all VSNs while BS-lectin is more highly expressed by fibers in the posterior AOB. Dashed boxes indicate the regions magnified for comparison in G-J. C-F) Resulting P-Values when comparing WT and Kirrel3 KOs when using various settings for block size (51, 101, 201) and minimum size limit (1, 20, 40, 60, 80, 100). The resulting p-values for average glomeruli size in the aAOB(C) and pAOB (D) and for the average number of glomeruli per section in the aAOB (E) and pAOB (F). Horizontal red dotted line indicates p-value = 0.05. G-J) Comparison of manual tracings of the WT (I,K) and Kirrel3 KO (J,L) glomeruli in the anterior (G, H) and posterior (I,J) AOB to the automated identification of glomeruli at block size 101 with minimum size limits at 1, 60, and 100. K-N) Automated quantifications at block size 101 for glomerular size (K,L) and # of glomeruli (M,N) without a minimum size limit (Limit # 1, K, M) small regions were included in the quantifications making any phenotypic differences across genotypes undetectable in both the anterior and posterior AOB. The same automated identification of glomeruli including a minimum size limit of 60 (L, N)) showed could detect differences in glomerular size (L) and (number (N) in the WT and Kirrel3 mutant mice in the pAOB, as expected. However, using these parameters our program also detected significant differences in glomerular size in the anterior AOB but no significant difference in the number of glomeruli.

To evaluate how the block size and minimum size limit would affect the detection of significant differences between the WT and Kirrel3 KOs we compared the resulting values obtained with different combinations of block size and minimum size limit through statistical analysis. The P-values from these analyses were graphed for glomerular size in the aAOB (Figure 4C) and pAOB (Figure 4D) as well as average number of glomeruli per section for the aAOB (Figure 4E) and pAOB (Figure 4F). We found that setting block size at value 101 selected most of the glomerular layer and allowed us to detect consistent statistical differences between genotypes in size (Figure 4C, D) and number of glomeruli (Figure 4E, F) in both anterior and posterior regions when minimum size limits were implemented.

To test the effects of various minimum size limits on detecting putative glomeruli across genotypes (compared with manually selected glomeruli in Figure 4G-J) we analyzed samples of Kirrel3 KOs and WT controls, implementing different size minimums, and then compared the average size and number of glomeruli for each condition. The results of these observations indicated that without setting a minimum size limit (Limit 1, Figure 4K, M) we could not detect significant differences in size (Figure 4K) and number (Figure L) of glomeruli between WT controls and Kirrel3 KOs. However, setting a minimum size limit of 60 allowed us to detect and confirm previously published differences between size (Figure 4M) and number of glomeruli per section (Figure 4N) in the pAOB (Prince *et al.* 2013). Notably, by setting limit sizes, and therefore creating specific cutoffs in the identification of the ROIs we also identified small but significant differences in the size (Figure 4L) in the aAOB which were not previously detected manually (Prince *et al.* 2013).

We found by testing these parameters that block size has a great influence on the regions recognized as glomeruli. In fact, by changing the block size we were able to obtain higher resolution in the identification of glomeruli as individual ROIs. By setting different values for block size (51, 101, 201 pixels) on the same biological samples we observed identified block 101 as the setting that yielded selections similar to manually made ROIs. These observations suggest that running comparative analysis with different block sizes could facilitate the identification of differences that can be further validated by human measurements (Figure 4).

Interestingly, using these parameters, we found that limiting the number of small artifacts detected by the program is essential for detecting statistically significant differences between phenotypes (Figure 4C-F). By analyzing samples from mutant mice that have been shown to have aberrant glomeruli size and number we observed that setting the minimum size for object recognition is necessary to detect significant differences across samples. In fact, when no minimum size limit was implemented the previously reported phenotypic differences when comparing Kirrel3 KO mutants and WT controls could not detected for glomeruli size (Figure 4K) or number (Figure 4M). However, by experimenting with different settings of block size and minimum size limits we found that in our experimental conditions that block size 101 and a minimum size limit of 60 were able to recapitulate the published differences between size (Figure 4L) and number of glomeruli (Figure 4N) in the pAOB between WT and Kirrel3 KOs (Prince *et al.* 2013). Notably, by using consistent parameters for glomeruli detection and setting minimum limit sizes, therefore creating specific cutoffs for the inclusion of identified ROIs, we also found subtle but significant differences in the size of the glomeruli in the aAOB (Figure 4M) that were not previously detected by manual quantifications (Figure 4C-D’’). These data, if further confirmed, could point to previously undetected effects of Kirrel3 mutation on neuronal populations.

In this article we present a new and free bioinformatic tool that allows us to perform automated quantification of structures, such as glomeruli in the AOB of rodents, using adaptive thresholding, which considers local differences in pixel intensity to identify regions that may be missed by using global thresholding, such as low intensity areas. Using this program, we were able to perform rapid systematic comparative analysis of AOB glomeruli size, number, and immunoreactivity between mice with different genetic makeup applying consistent parameters across an entire image set. In order to optimize glomeruli detection starting from digital images that vary in quality and signal-noise ratio we identified block size and minimum size limits as key parameters.

Optimization experiments for setting block size and minimum size limits should be performed for individual staining and sets of images as these could differ between labs for quality, contrast, definition, and noise. As in manual quantification, avoiding oversaturation of the images is fundamental and necessary for identification and selection of distinct regions. While the human eye may be able to infer the distinction between objects even in a slightly saturated setting based of other markers in the image, the program cannot make such distinctions as it uses adaptive thresholding to identify regions with similar intensity in a single channel (data not shown). Additionally, immunostaining and image acquisition should be performed as close to the same conditions and settings as possible.

Our program identifies putative glomeruli, using consistent parameters, where an ROI either define an entire glomerulus or a portion of it according to the adaptive thresholding (Figure 3D-D’’’, Figure 4G-J). Notably, though the identified shapes between human and automated quantifications appeared different the normalized results from each method yielded comparable statistical significances (Fig 3E-G). These data indicate that our system is sensitive enough to detect differences across genotypes and that using consistent parameters could facilitate the identification of subtle differences that can be lost due to time constraints and variability between investigators.

## ETHICS STATEMENT

All mouse studies were approved by the University at Albany Institutional Animal Care and Use Committee (IACUC).

## CONFLICT OF INTEREST

The authors declare that the research was conducted in the absence of any commercial or financial relationships that could be construed as a potential conflict of interest.

## FUNDING

Research reported in this publication was supported by: the Eunice Kennedy Shriver National Institute of Child Health and Human Development of the National Institutes of Health under the Awards R15-HD096411 (P.E.F), and R01-HD097331/HD/NICHD (P.E.F); the National Institute of Deafness and other Communication Disorders of the National Institutes of Health under the Award R01-DC017149 (P.E.F); National Institute on Drug Abuse of the National Institutes of Health under the Award R01 DA047410/DA/NIDA (A.K); UAlbany Next Research Frontier (NeRF) initiative (P.E.F and A.K); The Canadian Institutes for Health Research and the Natural Sciences and Engineering Research Council of Canada (J.-F.C).

## Pre-requisites

### 1 Installing Python 3.x

**Step 1:** Please visit official Python website (https://www.python.org).

**Step 2:** Download and install the latest Python 3.x installer corresponding to your operating system. (64-bit version is recommended.)

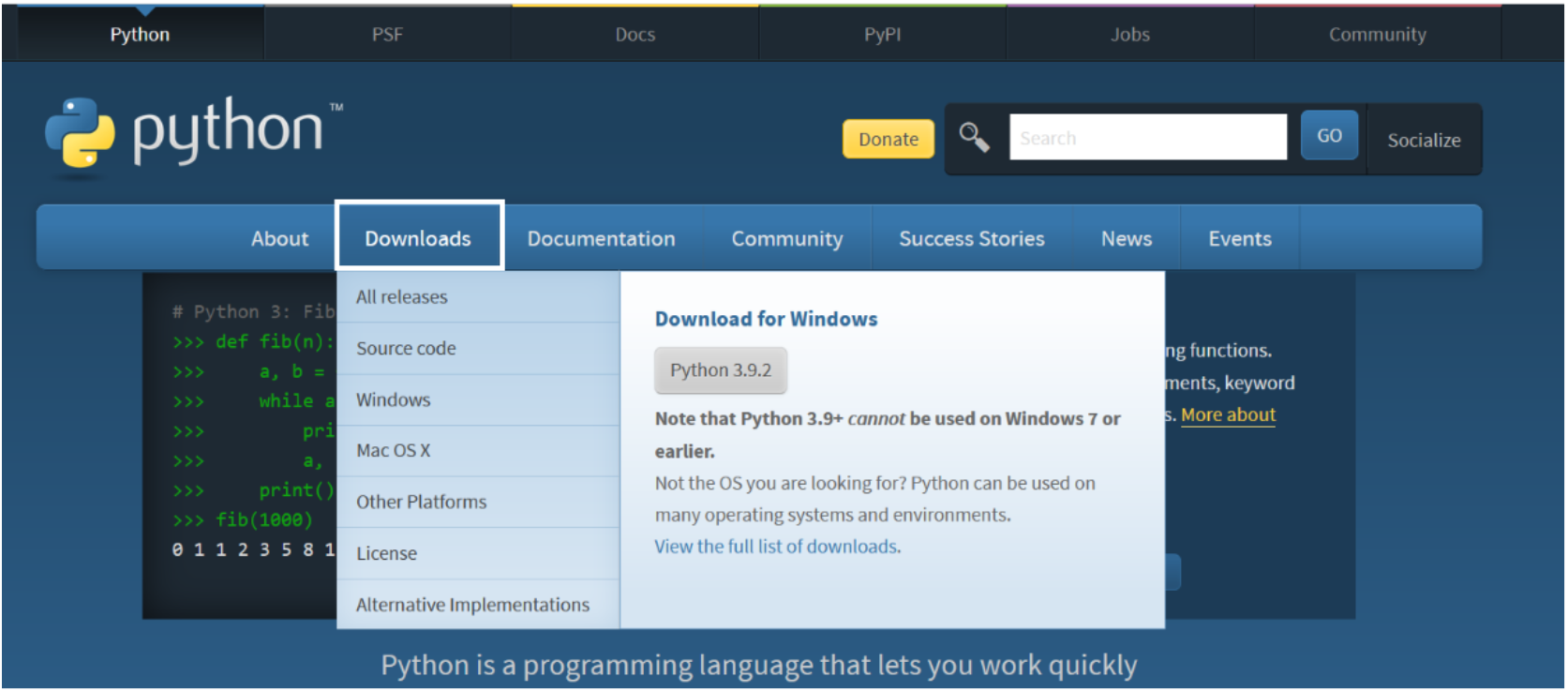

**Please note:** for windows users, please check mark “Add Python 3.x to Path” on installation.

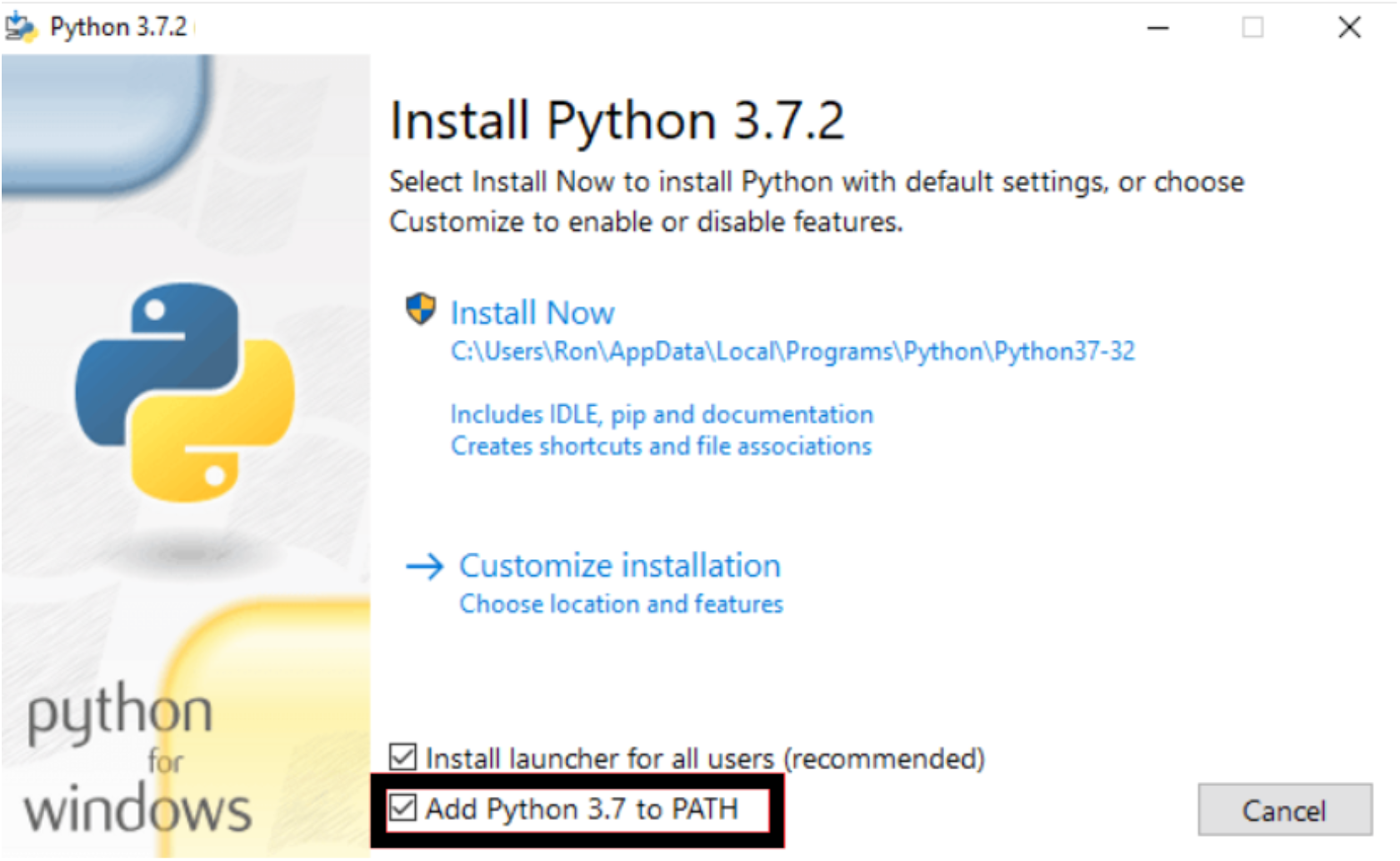

**Step 3:** check if Python is installed properly. Please open command prompt CMD (or terminal for MAC/Linux users) and enter the following command: **python --version**

The result should be as follows: **Python 3.7.2**

**Please note:** if you got different version of Python on CMD/Terminal output, that means you have installed multiple Python engine and the default is the one that appeared in CMD/Terminal. It is strongly recommended to uninstall all existing Python engines and reinstall the desired one again.

### 2 Installing required Python packages

All required packages to run the software properly are listed into a file called “requirements.txt”. This file should be available on repository.

**a)** In order to install packages, please visit “<u>Glomeruli-Analyzer</u>” project webpage download it as a zip file.

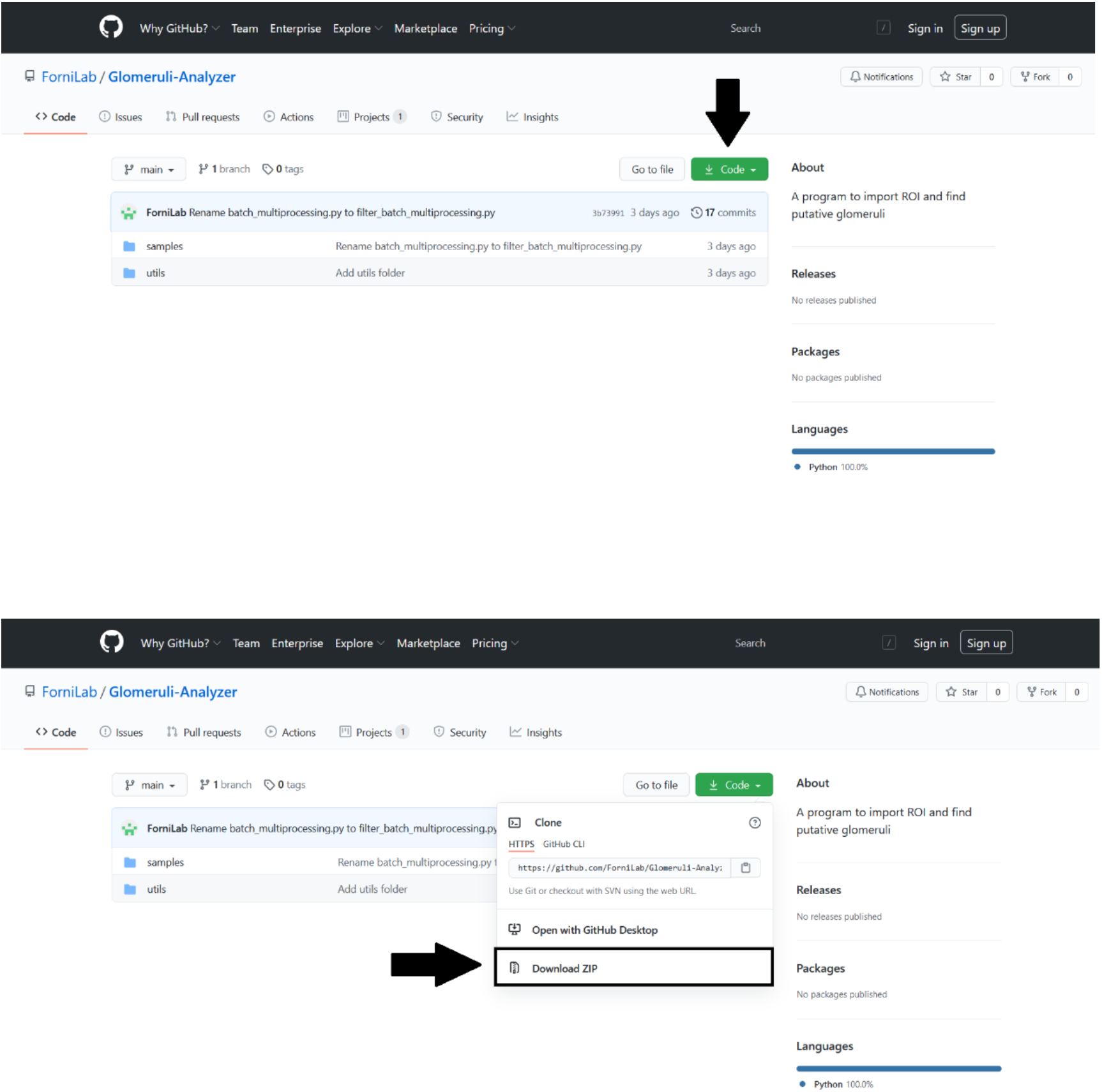

**b)** Please extract downloaded zip file in a folder and follow navigate to this path to get “requirements.txt”:

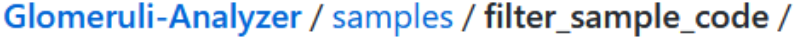

Next, open CMD/Terminal in the corresponding folder that you have put the downloaded file and enter this code into CMD/Terminal: **pip3 install -r requirements.txt**

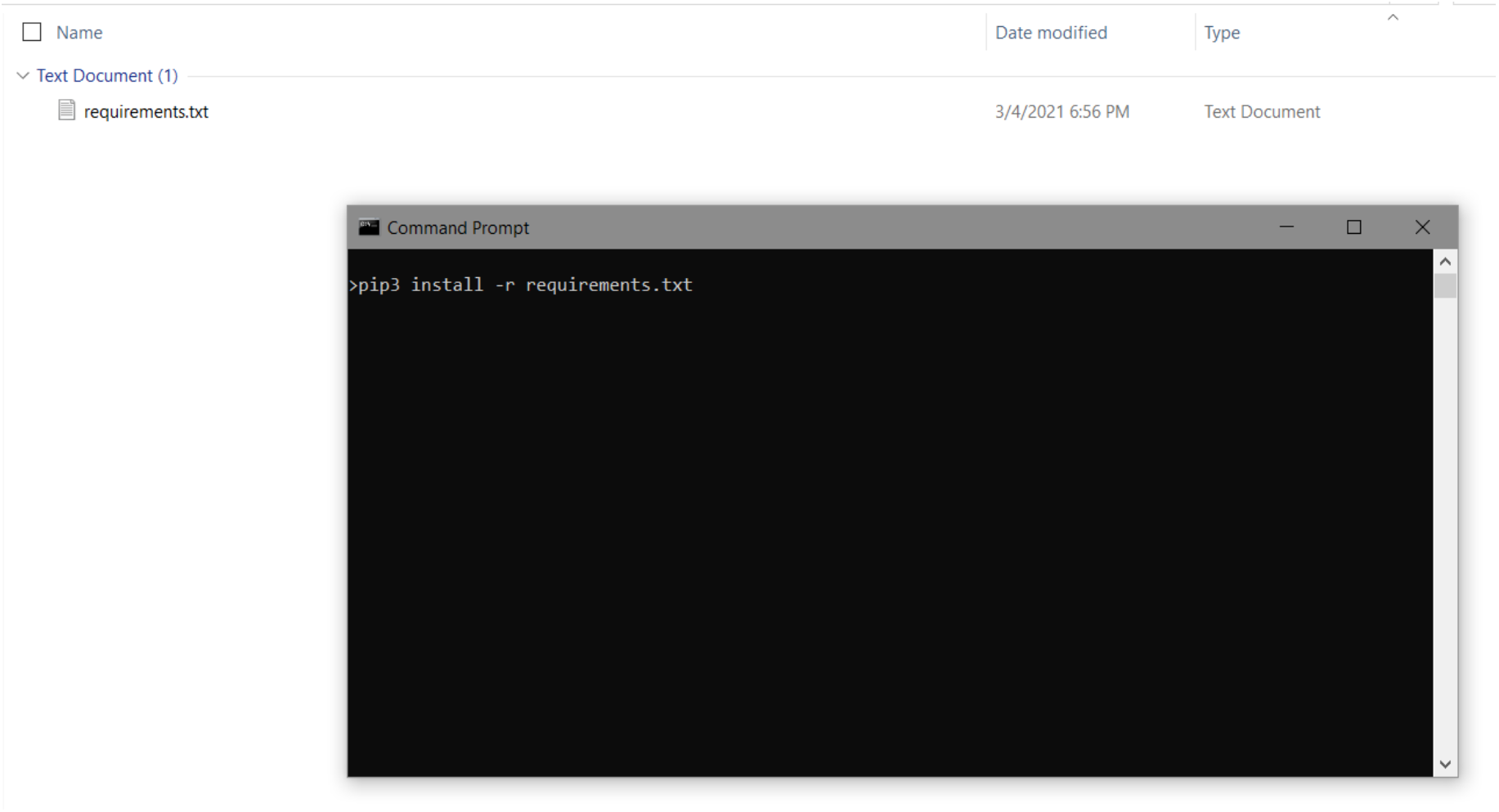

The required packages should start to install afterwards. Otherwise please check what packages are not able to be downloaded/installed and look for the error through pythons forums to throubleshoot it.

If everything installed properly and you run the code again should should receive the following message in CMD/Terminal: **Requirement already satisfied: …**

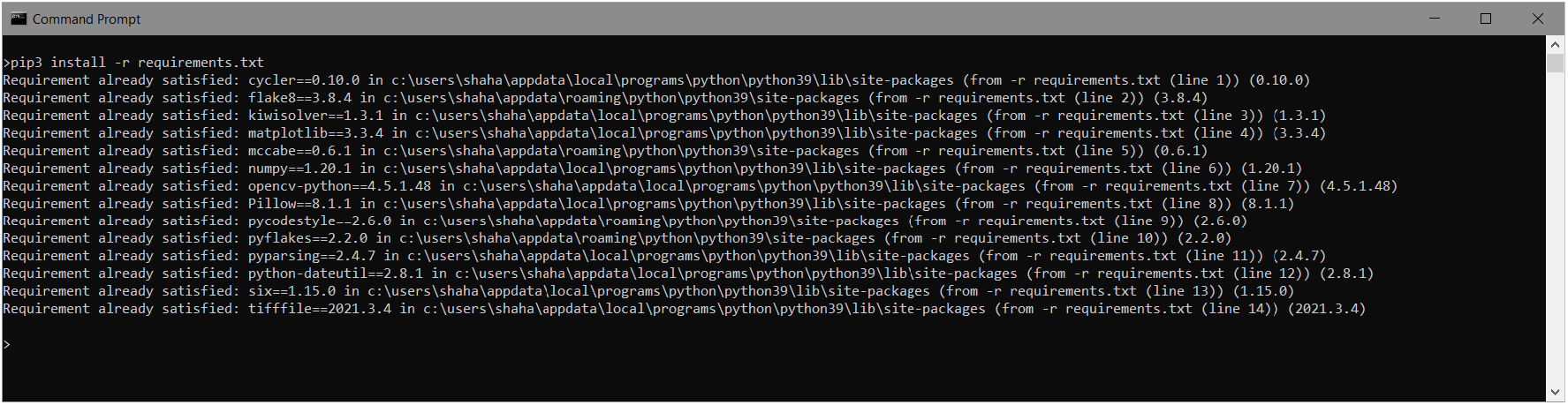

## How to run the program

There are various ways to run the program in different operating systems. In general, we divide it into two different methods:

1- Run by CMD/Terminal (multiprocessing)

Please download the whole folder as a zip file and extract it into a folder in your local machine.

**Step1:** put all images into “images” folder. (subfolders would be ignored)
**Step2:** change setting in “setting_batch.py” into your desired values.
**Step3:** run CMD/Terminal in the extracted folder and enter the following code:

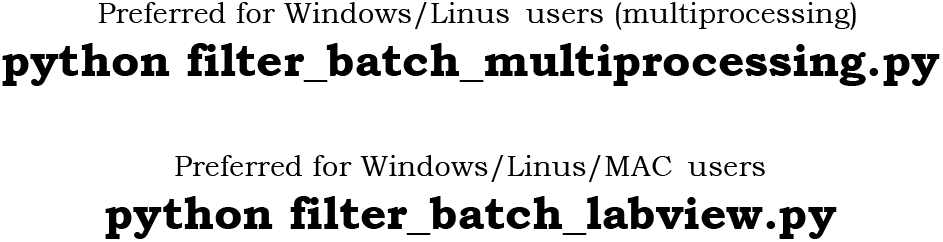 Please wait until the process is completed.
**Step4:** check “output” folder for all analyzed images output. Each images analysis should be separated into folders.
**Step5:** there is an “output.txt” in “output” folder which include all parameters for all images. One may copy all data and paste it into an excel file.

2- Using compiled GUI or LabVIEW subVI

In order to use compiled GUI please choose the corresponding file for your operating system and run the compiled standalone program.

a) **(Recommended)** To use LabVIEW sub-VI:

i. Please download, install, and activate LabVIEW community (above 2020) from its official website: (https://www.ni.com/en-us/support/downloads/software-products/download.labview.html) **Please note:** if you installed Python 3.x 64-bit you have to install LabVIEW 2020 64-bit, and vice versa for 32-bit version as well.
ii. Open GUI.vi in LabVIEW and run it.

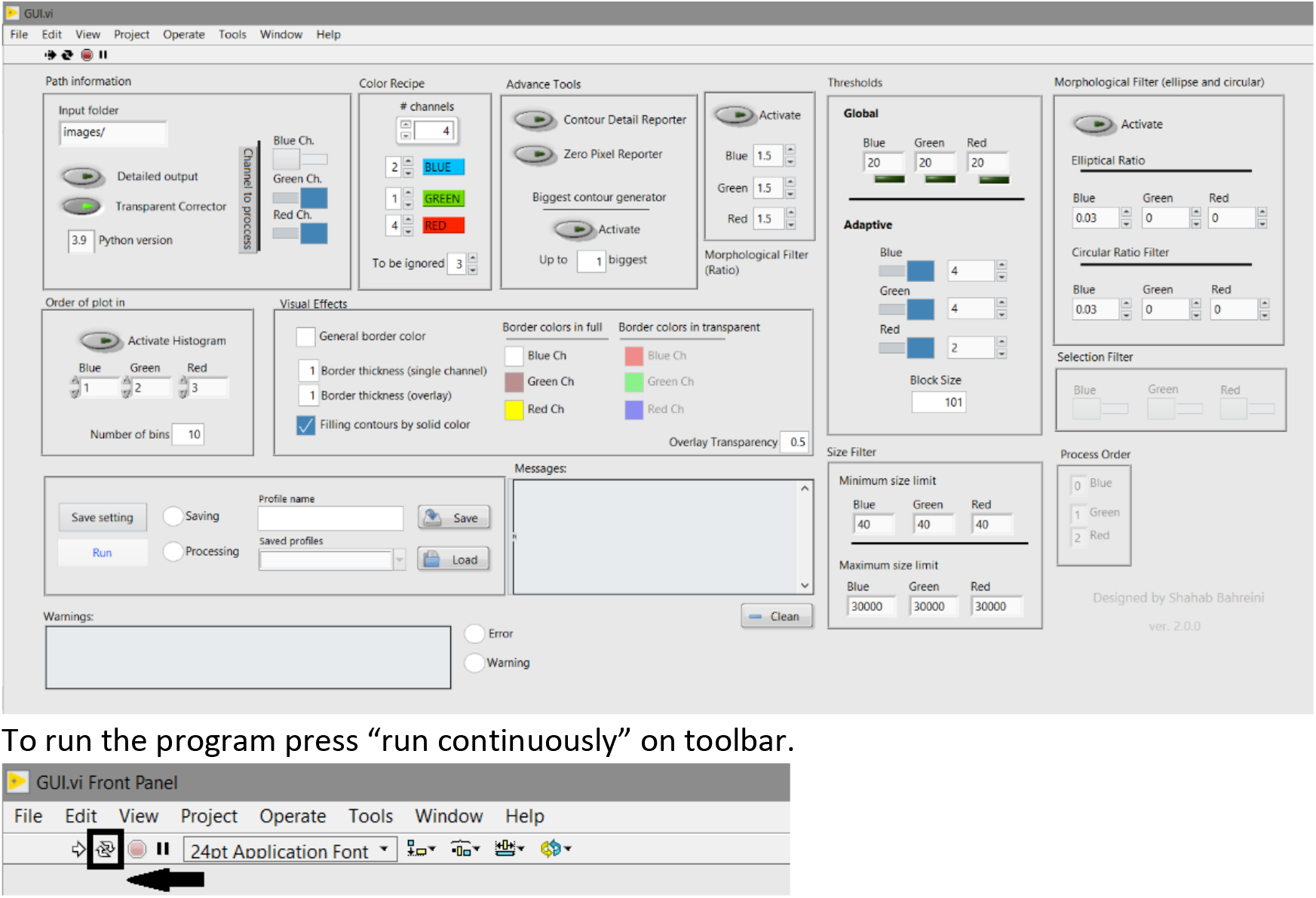

b) Run compiled version:

<u>Windows OS:</u> please run “Application.exe” and on the very left of the toolbar click “run continuously”.

<u>MAC OS:</u> please run “Application-MAC” and on the very left of the toolbar click “run continuously”.

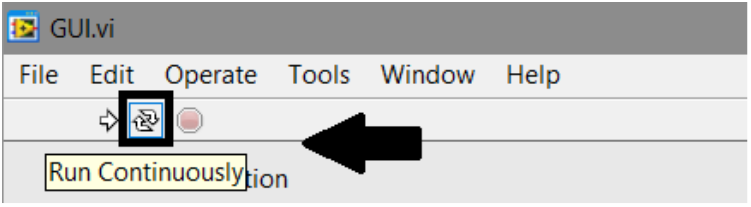

To process an RIO, you may choose your best setting by buttons and available controls and after all parameters are set, please click “Save setting”:

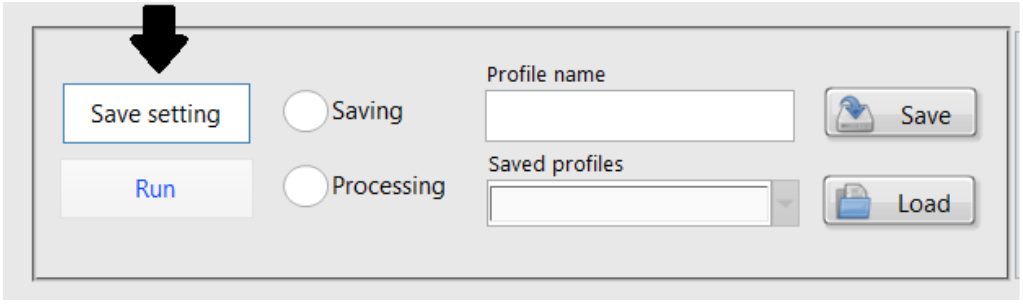

The following message should be appeared in message box. And in the program root folder “setting_batch.py” should be added/created at the same time.

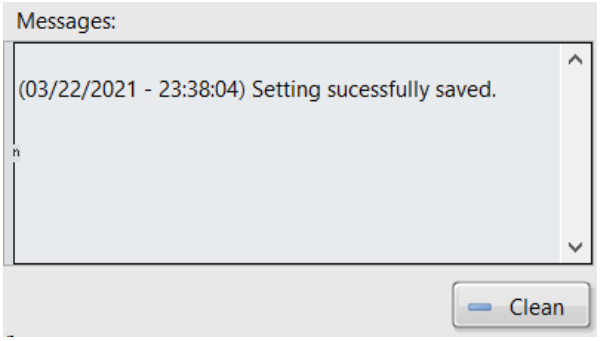

**Please note** if “setting_batch.py” is an old one (check the modified time of the file), that means GUI did not run properly. Then please use another mentioned (1 or 2) method.

After saving setting, by clicking on “Run” button “outputs” folder should be updated with the last process images and the following message should be appeared there.

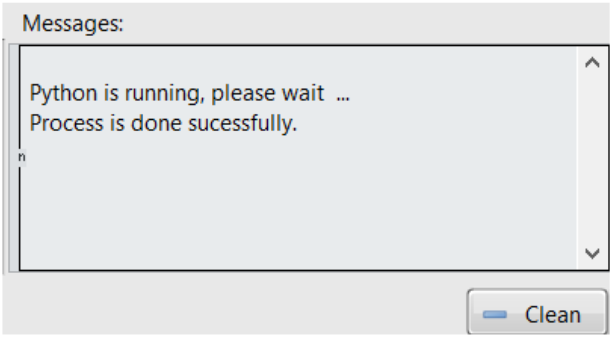

**Please note** if “outputs” folder did not update with new processed file, that means GUI did not run properly and it is highly recommended to run the program by Terminal/CMD. To do so, please open Terminal/CMD in the program root folder and run the following code:

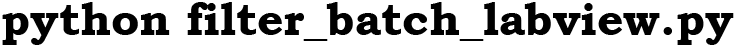

In terminal you would see, total process time, number of files and some process information per each image as follows. Below is the result of processing red and green channel for “test.png” sample file which is available in folder “images”.

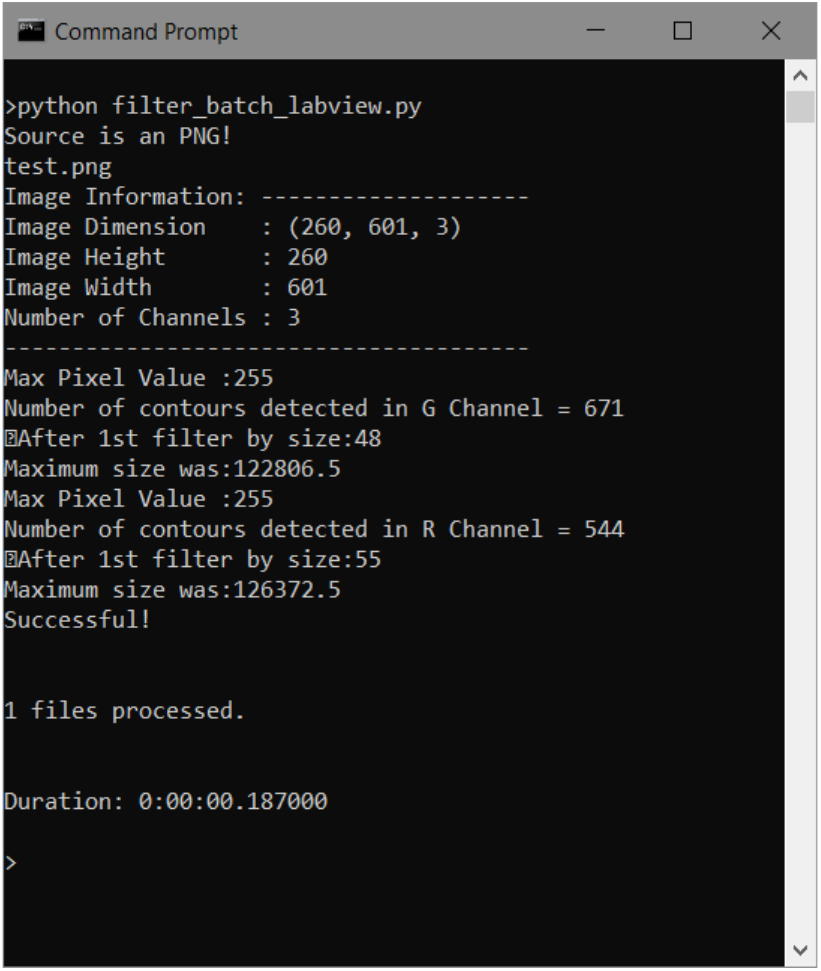

## Filter tools

The goal of using morphological filters is to ignore axon like identified shapes or in other words set a limit of acceptance of elongated contours.

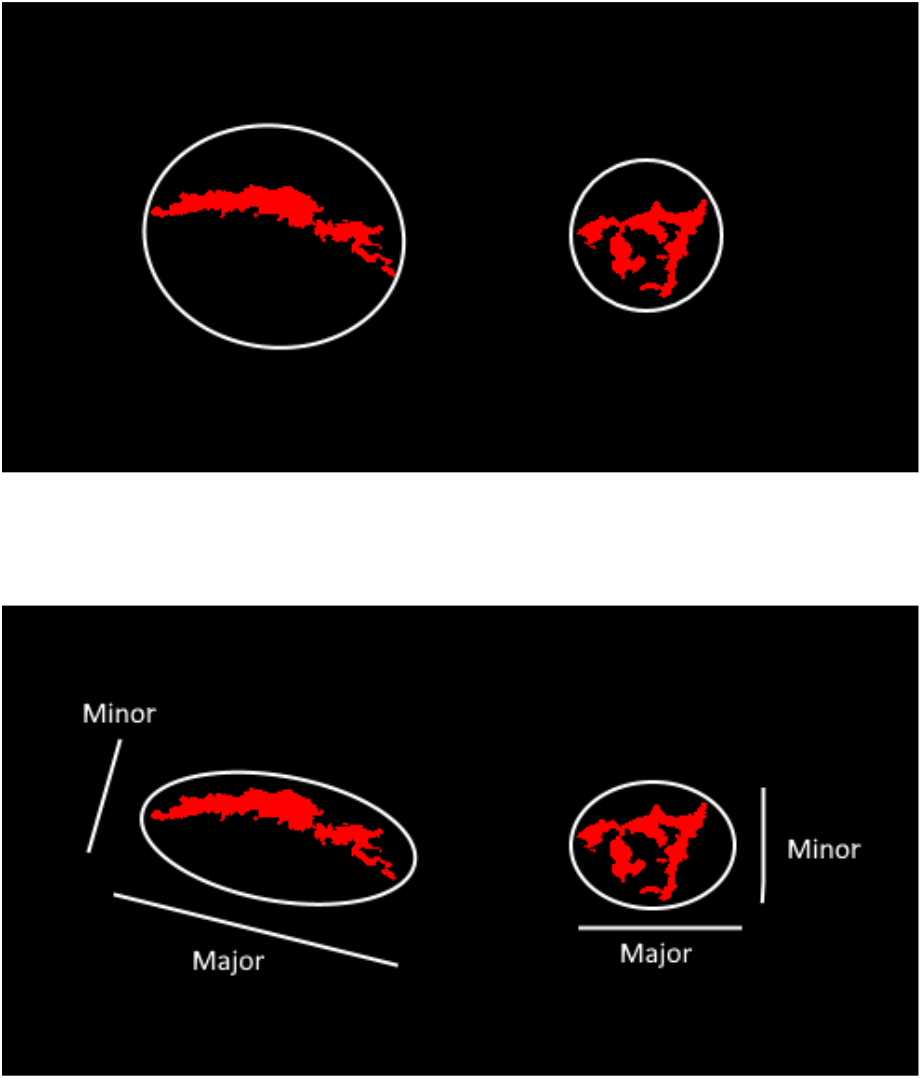

In <u>circular filter</u>, program encircles contours into a circle. By calculating the contour area and circle area the following ratio is determined: *Size*/πr^2^

This ratio indicates how thin and elongated is the contour. So, for axon like contours this ratio is low. Circular filter tries to put a limit of acceptance for this ratio to ignore axon like contours.

<u>Ellipse filter</u> inherited the same method of circular filter; however, it encircles contours with an ellipse. The *minor axes*/*major axes*. is calculated to check how elongated is the contour. Smaller ratio means more elongated shape.

In both cases acceptable range of circular/ellipse ratio is defined as below:

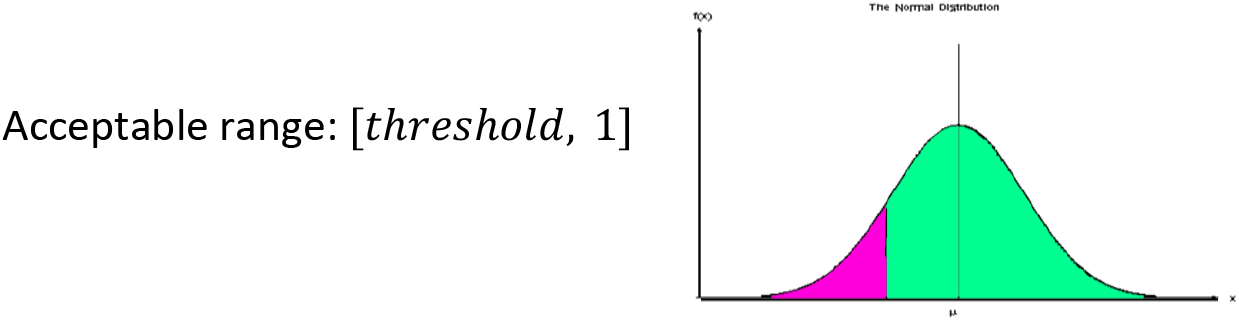

<u>Ratio filter</u> also is another tool to help removing axon like shapes with a different method. In this method program computes the following ratio of each contour: *primeter*/*area*

The larger ratio indicates the more snakier shapes (or contours with more thin wire like branches). To set a limit of acceptance, by setting a positive number (λ), program ignores all contours that has a ratio below the following range:

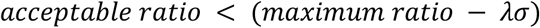

Here σ indicates standard deviation of ratio distribution.

## Histogram

In order to determine threshold ratio in morphological filters, histogram of ratio distribution helps. This feature allow program to generate histogram of ratio distribution of circular and ellipse filters. Below is a sample of generated histogram for RGB channels (see figure 1,2).

**Figure 1.**
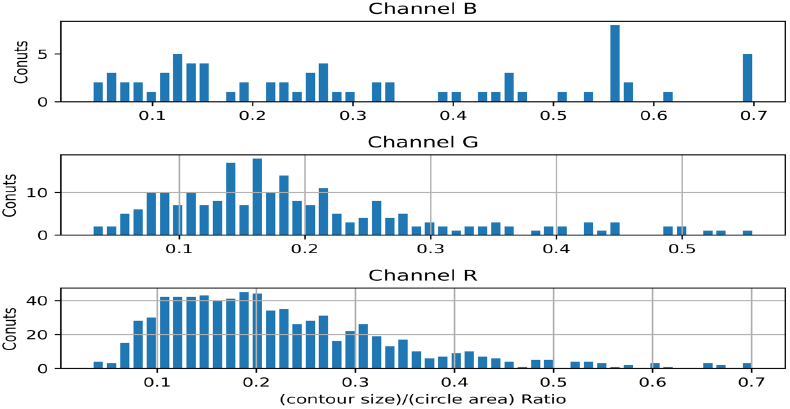
Circular ratio distribution histogram

**Figure 2.**
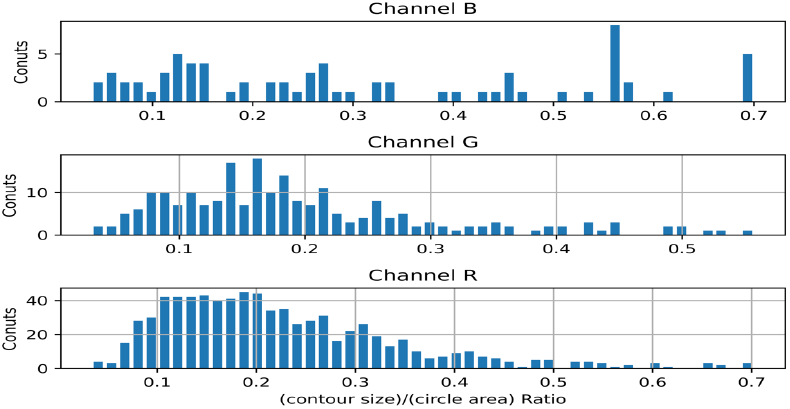
Ellipse ratio distribution histogram

## Size limit filter

The goal of this tool is to eliminate miniscule identified contours. Figure 3 is a sample of processed raw ROI image which contains a large number of tiny patterns.

**Figure 3.**
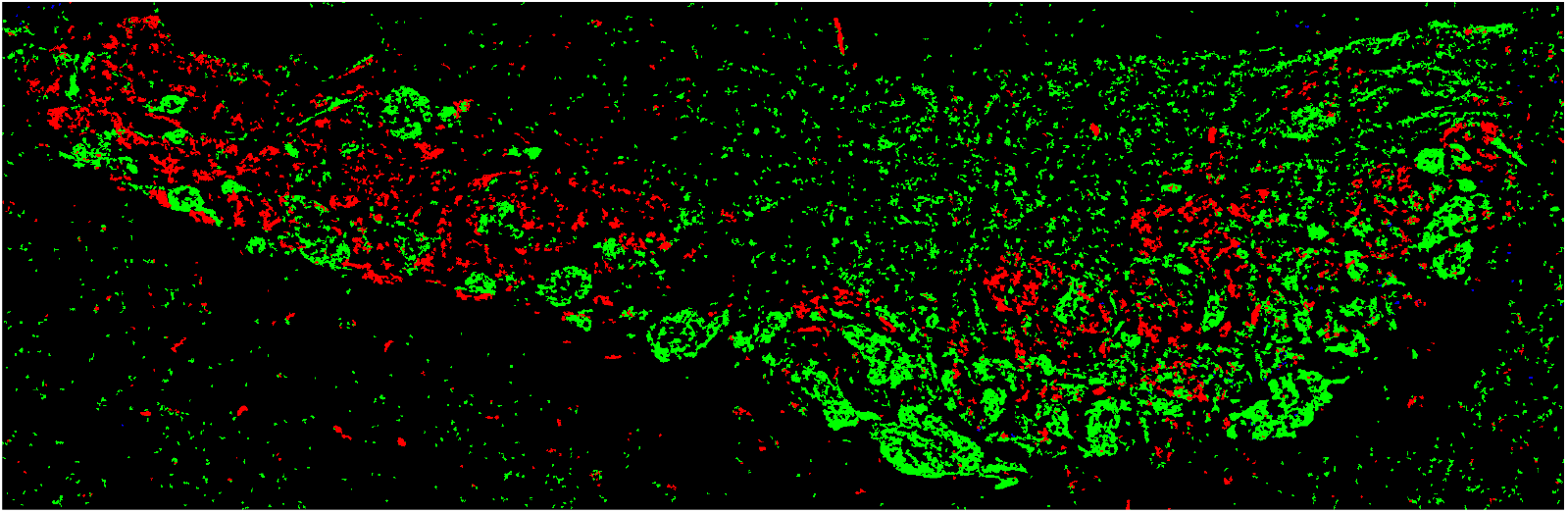
sample of output for a raw ROI. Too many tiny patterns are identified in image.

However, set a size limit threshold helps to purify program output (see Figure 4).

**Figure 4.**
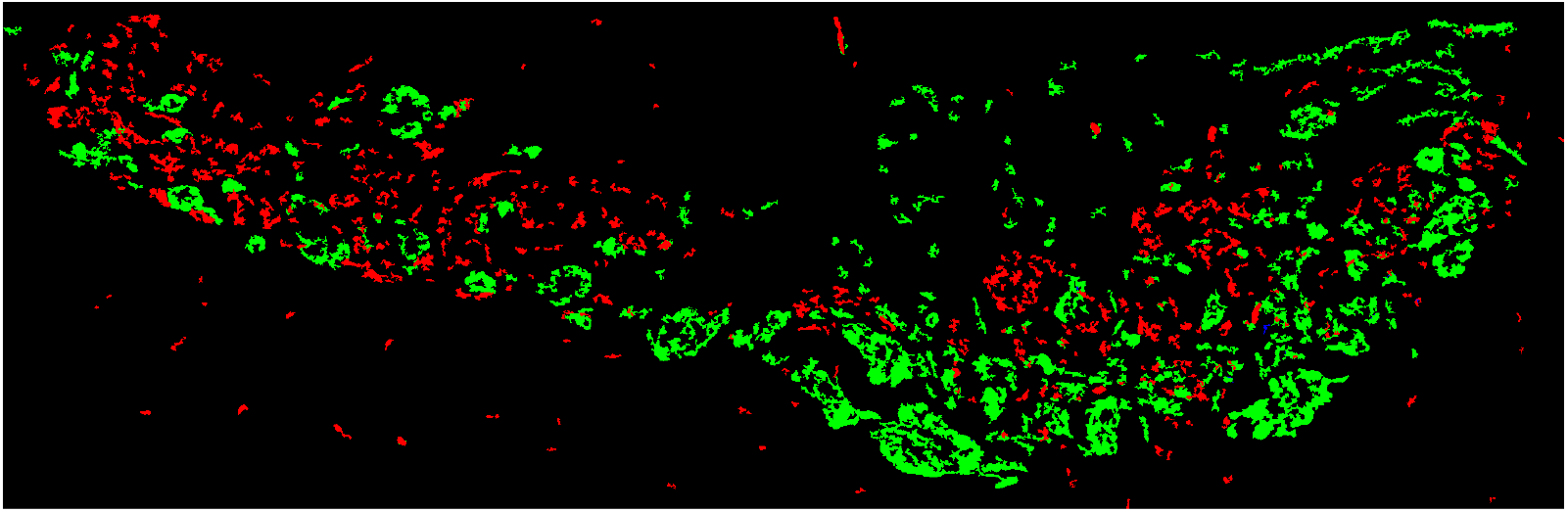
Set size limit 10, 40 and 10 for RGB channels respectively helps to eliminate small redundant patterns.

## Technical Setting Description

Path Information

Path Information
| Name | Description | Value | Available in GUI<br>(graphical interface) | Available in file<br>(manual edit) |
| --- | --- | --- | --- | --- |
| Input folder | Folder name which contains ROIs. (no path is acceptable, just the name + "/" ) | Name + "/" | ✓ | ✓ |
| Detailed output | Enable/Disable contour details per images. This option allows program to generate full detailed result per ROI. | True/False | ✓ | ✓ |
| Transparent corrector | Removes any available solid background from ROIs. | True/False | ✓ | ✓ |
| Channel process | Determines which channel should be processed. | True/False | ✓ | ✓ |

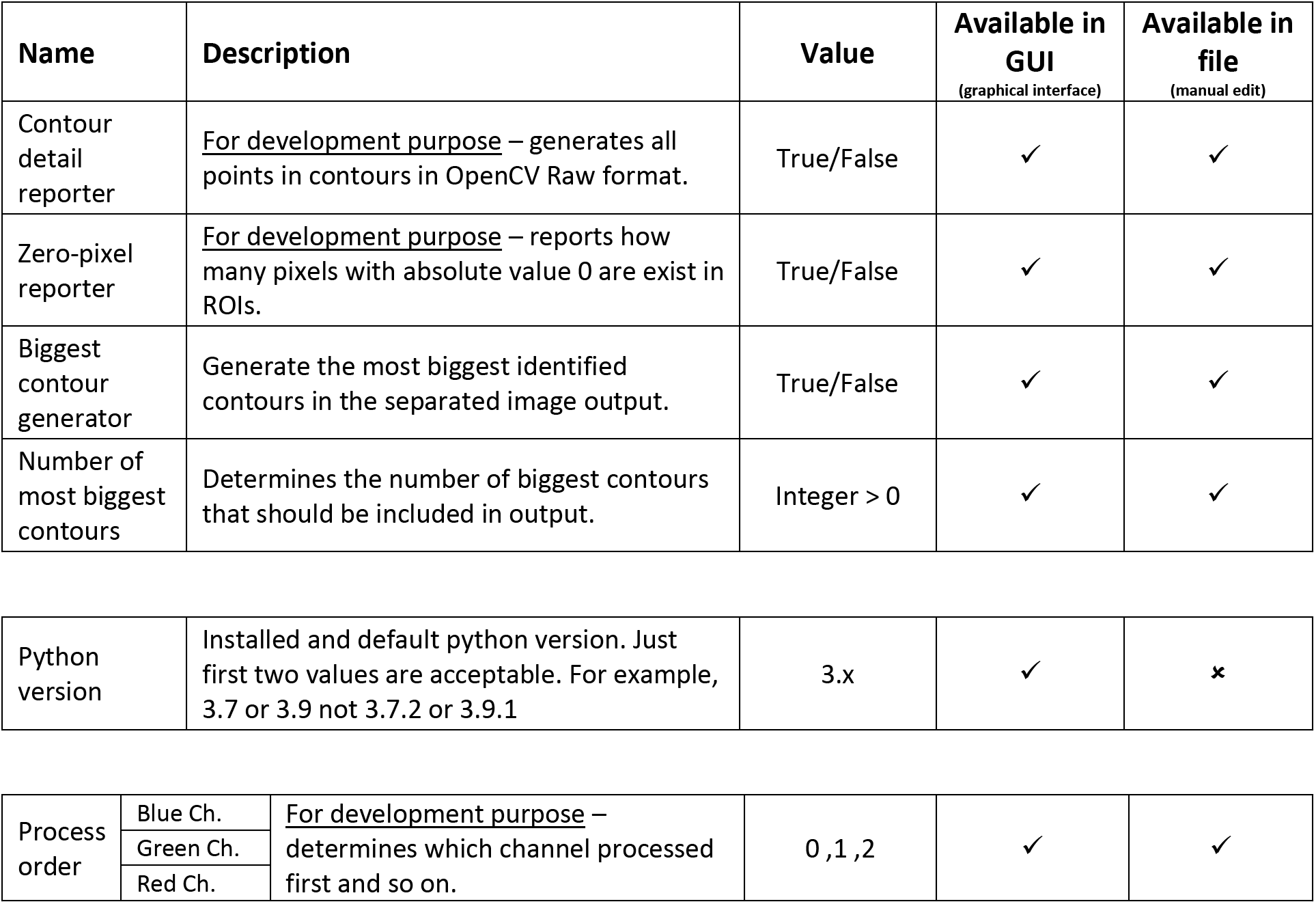
Advanced Tools

Advanced Tools
| Name | Description | Value | Available in GUI<br>(graphical interface) | Available in file<br>(manual edit) |
| --- | --- | --- | --- | --- |
| Contour detail reporter | <u>For development purpose</u> – generates all points in contours in OpenCV Raw format. | True/False | ✓ | ✓ |
| Zero-pixel reporter | <u>For development purpose</u> – reports how many pixels with absolute value 0 are exist in ROIs. | True/False | ✓ | ✓ |
| Biggest contour generator | Generate the most biggest identified contours in the separated image output. | True/False | ✓ | ✓ |
| Number of most biggest contours | Determines the number of biggest contours that should be included in output. | Integer > 0 | ✓ | ✓ |
| Python version | Installed and default python version. Just first two values are acceptable. For example, 3.7 or 3.9 not 3.7.2 or 3.9.1 | 3.x | ✓ | ✗ |

|  |  |  |  |  |  |
| --- | --- | --- | --- | --- | --- |
| Process order | Blue Ch. | <u>For development purpose</u> – determines which channel processed first and so on. | 0 ,1 ,2 | ✓ | ✓ |
|  | Green Ch. |  |  |  |  |
|  | Red Ch. |  |  |  |  |

Thresholds

Thresholds
| Name |  | Description | Value | Available in GUI<br>(graphical interface) | Available in file<br>(manual edit) |
| --- | --- | --- | --- | --- | --- |
| Global | Blue | Activates when adaptive threshold is disabled. A small LED indicator is below each channel box which turns on when global threshold activates for each channel. | Integer > 0 | ✓ | ✓ |
|  | Green |  |  |  |  |
|  | Red |  |  |  |  |
| Adaptive | Blue | Enable/Disable adaptive threshold per channel. | Integer > 0 | ✓ | ✓ |
|  | Green |  |  |  |  |
|  | Red |  |  |  |  |
| Block size |  | Determines adaptive threshold block size | Odd Integer > 8 | ✓ | ✓ |

Morphological Filter (ellipse and circular)

Morphological Filter (ellipse and circular)
| Name |  | Description | Value | Available in GUI<br>(graphical interface) | Available in file<br>(manual edit) |
| --- | --- | --- | --- | --- | --- |
| Activate |  | Enable/Disable both ellipse and circular filters | True/False | ✓ | ✓ |
| Elliptical Ratio Filter | Blue | This filter tries to encircle each contour by an ellipse. It determines the ratio of (minor axis)/(major axis). Next, filters out all those contours which has smaller value than computed ratio. | > 0 | ✓ | ✓ |
|  | Green |  |  |  |  |
|  | Red |  |  |  |  |
| Circular Ratio Filter | Blue | This filter tries to encircle each contour by a circle. For circular filter it computes ratio of (contour area)/(circle area). Next, filters out all those contours which has smaller value than computed ratio. | > 0 | ✓ | ✓ |
|  | Green |  |  |  |  |
|  | Red |  |  |  |  |

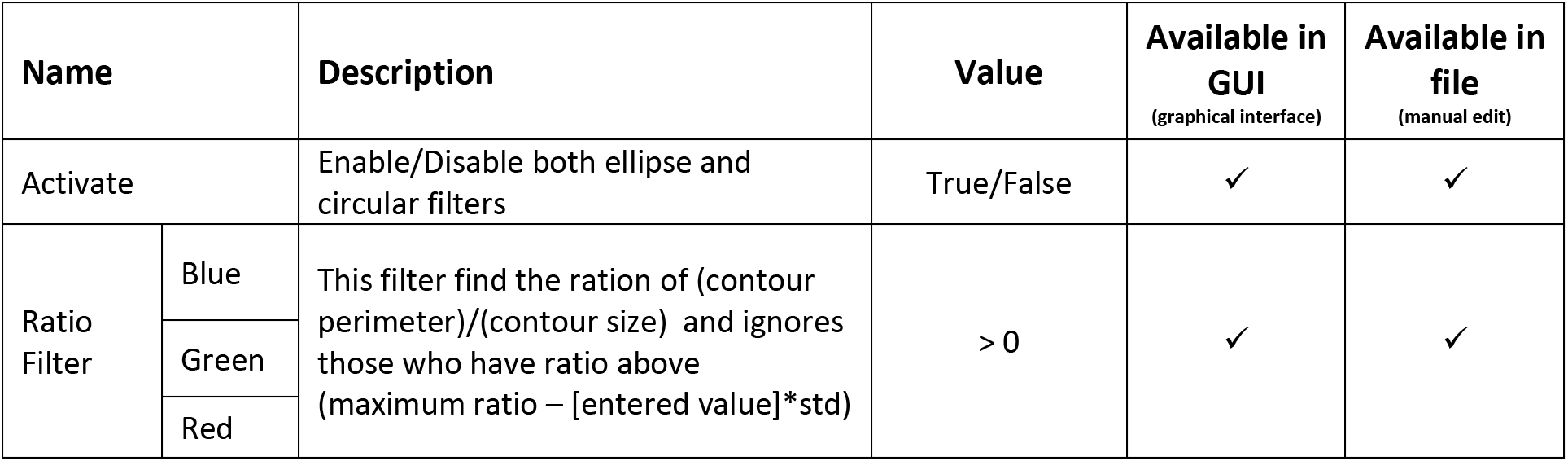
Morphological Filter (Ratio)

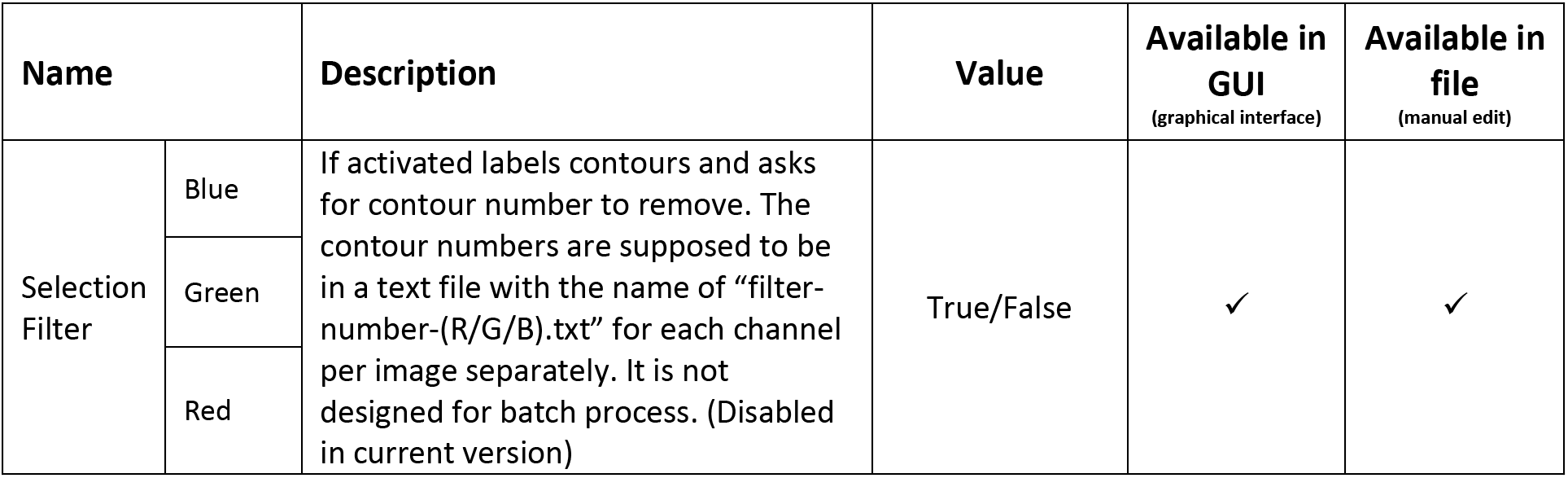
Selection Filter

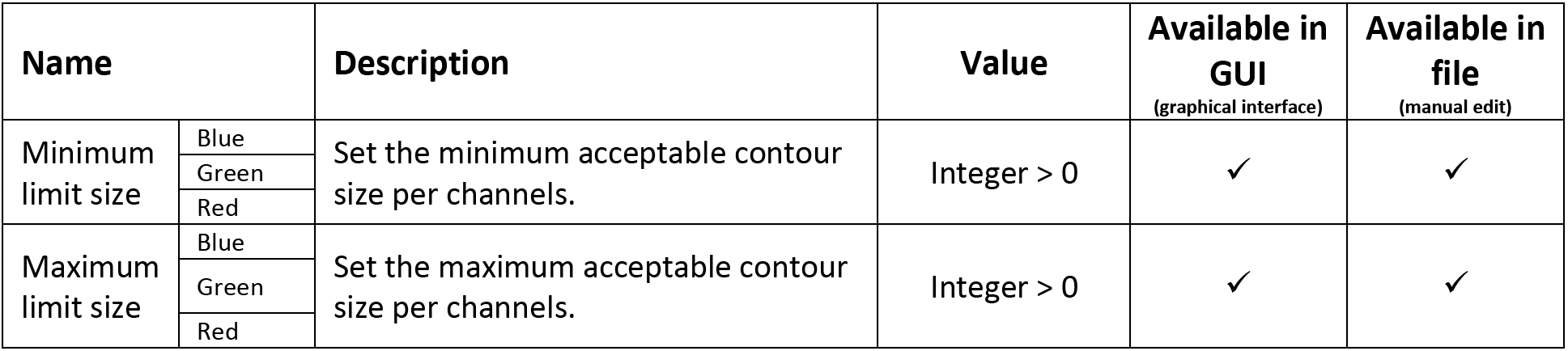
Size Filter

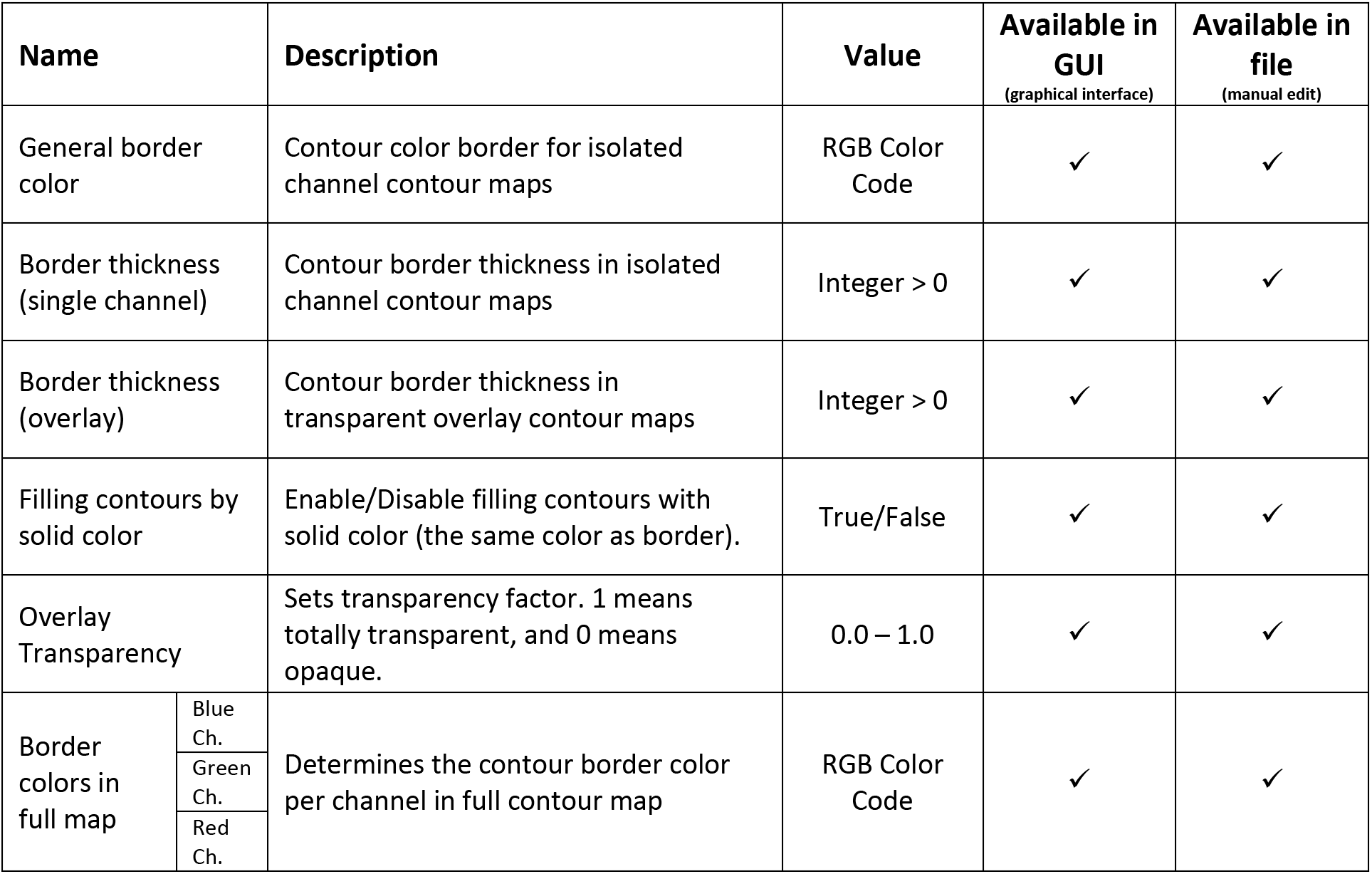
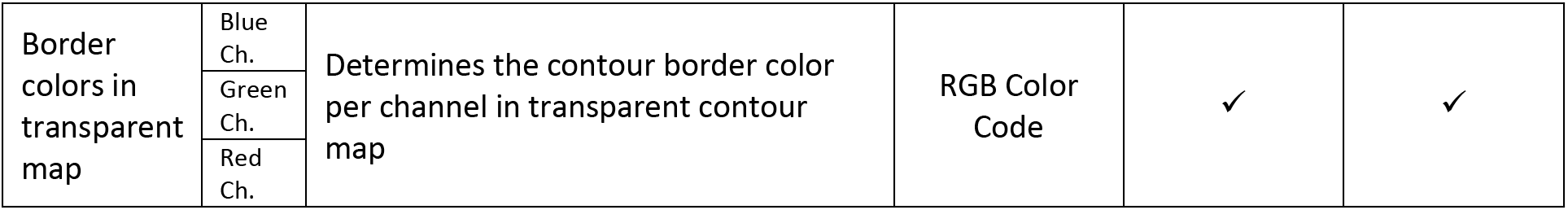
Visual Effects

Order of plot in histogram

Order of plot in histogram
| Name |  | Description | Value | Available in GUI<br>(graphical interface) | Available in file<br>(manual edit) |
| --- | --- | --- | --- | --- | --- |
| Activate Histogram |  | Enable/Disable generating histogram for elliptical and circular filter ratios. | True/False | ✓ | ✓ |
| Channel Order | Blue | Order channels for generated histogram. 1 has the most priority. | 1, 2, 3 | ✓ | ✓ |
|  | Green |  |  |  |  |
|  | Red |  |  |  |  |
| Number of bins |  | Determines the number of bins for generating histogram. | Integer > 0 | ✓ | ✓ |

Color Recipe

Color Recipe
| Name | Description | Value | Available in GUI<br>(graphical interface) | Available in file<br>(manual edit) |
| --- | --- | --- | --- | --- |
| # channels | Determines how many channels are integrated into ROI. (In this version, program just accept 3 or 4) | 3, 4 | ✓ | ✓ |
| BLUE | The channel number to set as blue channel. Maximum possible value is the ROI channel number. | Integers < # channels | ✓ | ✓ |
| GREEN | The channel number to set as green channel. Maximum possible value is the ROI channel number. | Integers < # channels | ✓ | ✓ |
| RED | The channel number to set as red channel. Maximum possible value is the ROI channel number. | Integers < # channels | ✓ | ✓ |
| To be ignored | Channel number which is supposed to be ignored in analysis. | Integers < # channels | ✓ | ✓ |

Save & Run

Save & Run
| Name | Description | Value | Available in GUI<br>(graphical interface) | Available in file<br>(manual edit) |
| --- | --- | --- | --- | --- |
| Save setting | Saves the current setting as the one which program use it. | - | ✓ | ✓ |
| Run | Run the process. Please wait until processing LED turns off. Program runs with the last saved setting. | - | ✓ | ✓ |
| Profile name | The name of current setting that wants to be saved and be loaded in future. | Integers < # channels | ✓ | ✓ |
| Save | Saves the current values into a file which can be loaded in future. All profiles will save into “profiles” folder with “.ini” file extension. | Integers < # channels | ✓ | ✓ |
| Saved profiles | This drop-down list shows all available profile to choose and load. | Integers < # channels | ✓ | ✓ |
| Load | Loads the selected profile from drop-down list. |  |  |  |

## Notes

### Competing Interest Statement

The authors have declared no competing interest.

